# The Polyploid History of Cultivated Dahlia

**DOI:** 10.64898/2026.07.24.740608

**Authors:** Zachary Meharg, Kristine Albrecht, Jenell Webber, Lori Beth Boston, Jane Grimwood, Virginia Walbot, Alex Harkess

## Abstract

The cultivated dahlia has a complex evolutionary history that is likely the driver for its diverse floral forms. Previous studies have proposed cultivated dahlia as an allotetraploid, autotetraploid, and in one study an auto-allooctoploid. Here, we used PacBio HiFi long-reads and Dovetail Omni-C reads to assemble and annotate a genome for a tetraploid *Dahlia variabilis* cultivar Edna C. We identified ‘Edna C’ as a segmental tetraploid with signature of auto- and allo-polyploidy events based on subgenome-specific repeat partitioning and single-copy gene tree construction within homoeologous groups. Additionally, we identified a cultivated dahlia*-*specific centromeric monomer shared between *Dahlia variabilis* ‘Edna C’ and the *Dahlia pinnata* ‘Kelvin Floodlight’. Based on finding three major chromosome clusters, we propose that *Dahlia variabilis* ‘Edna C’ is a segmental tetraploid with evidence of auto- and allopolyploidy.

## Introduction

Cultivated dahlia (*Dahlia variabilis* Willdenow, *Dahlia* x *pinnata* Cavanilles, *Dahlia* x *cultorum* Thorsud & Reisater, or *Dahlia* x *hortensis* Guillaumin) is a perennial herbaceous ornamental species complex that has gone through several taxonomic classifications. Cultivated dahlia is a member of the *Dahlia* genus located within the Asteraceae family, which contains more than 10% of all flowering plant species (Mandel 2019). The *Dahlia* genus is comprised of both cultivated and wild dahlias totaling 43 described species, with a center of origin in Central Mexico (Reyes-Santiago et al. 2024). The first observation and record of wild dahlias was published in 1552 in *The Bandianus Manuscript* (Sorensen 1970). Dahlias were then introduced into Spain around the late 18^th^ century, where they were propagated and dispersed to other European countries through the Royal Botanical Garden of Madrid. Over the next 300 years, an explosion of cultivars was introduced around the world, leading to the widespread ornamental cultivation of dahlias. Cultivated dahlias are known for their diversity of blooms, with more than 50,000 cultivars registered with the American Dahlia Society (American Dahlia Society, 2026). This diversity encompasses floral forms ranging from an abundance of male-sterile ray florets, the elongation and pigmentation of discs, the doubling and elongation of ray florets, the conversion of reproductive structures into petals within ray florets, and blooms spanning the color spectrum (with the exception of blue and green). This phenotypic diversity, along with its relatedness to other agriculturally important Asteraceae such as sunflower (*Helianthus annuus*), lettuce (*Lactuca sativa*), and artichoke (*Cynara scolymus*), positions cultivated dahlia as a model system for exploring crop-relevant mutations in floral architecture.

Cultivated dahlias are polyploid relative to wild species in the genus, but it remains unclear if they are derived from auto-, allo-, or a segmental polyploidy event (Lawrence 1931, Gatt 2008, Schie 2014). Previous studies have sought to address *Dahlia* evolution using historical records, biochemistry, and ITS marker tree construction (Sorensen 1970; Ginannasi 1975). The historical records of *Dahlia* discuss a “double-bloomed” plant being grown in botanical gardens in Europe and speculate that it was a hybrid between *Dahlia coccinea* and *Dahlia sorensenii*, based on the respective floral pigmentation of each species: red, orange, and yellow for *D. coccinea,* and pink and lavender flowers for *D. sorensenii* (Dean 1903). Through traditional Mendelian breeding experiments, it was identified that floral color phenotypes were controlled by anthocyanins and flavones (Lawrence 1929). This was supported through biochemical analysis of cultivated and wild *Dahlia*, showing the *Dahlia* genus is composed of two distinct groups based on the presence or absence of anthocyanins and flavones (Ginannasi 1975); because cultivated dahlias contained both, this supported the hypothesis that they are a hybrid between the two groups within the genus. Nonetheless, it has been difficult to confidently identify the progenitors of the present-day cultivated dahlias because there is a lack of high-resolution genetic data for the majority of the *Dahlia* genus, and the unresolved polyploid history within the genus (Saar 2003, Gatt 2000, Gatt 1999, Gatt 2000, Schie 2013). A genome for cultivated dahlia, *Dahlia pinnata* ‘Kelvin Floodlight’, was recently published, presenting an autotetraploid genome containing 16 homoeologous groups with 4 chromosomes each (Wang 2024). Given the diversity within the genus, it is unclear if this genome is the same species as *Dahlia variabilis,* and if this genome architecture and type of polyploidy is a conserved feature among all cultivated dahlias in the genus.

To address these questions, we generated a chromosome-scale genome assembly for a tetraploid *D. variabilis* cultivar, Edna C (2n = 4x = 64). This cultivar was selected to be the genome line through a collaboration with the American Dahlia Society (ADS), a horticultural society founded in 1915 with membership spanning the everyday gardener to large-scale commercial dahlia growers. ‘Edna C’ was chosen as the ADS “Best Dahlia of the Past 50 Years” reflecting its longevity and numerous wins in the ADS judging and competition circuit (Figure 1a). Using ‘Edna C’ as the reference, we identified the genome structure of a cultivated dahlia, investigated sub-genome dynamics across 64 chromosomes with several complementary methods, and identified a centromere motif in cultivated dahlia that is shared with the recently published ‘Kelvin Floodlight’ genome.

**Figure 1:**
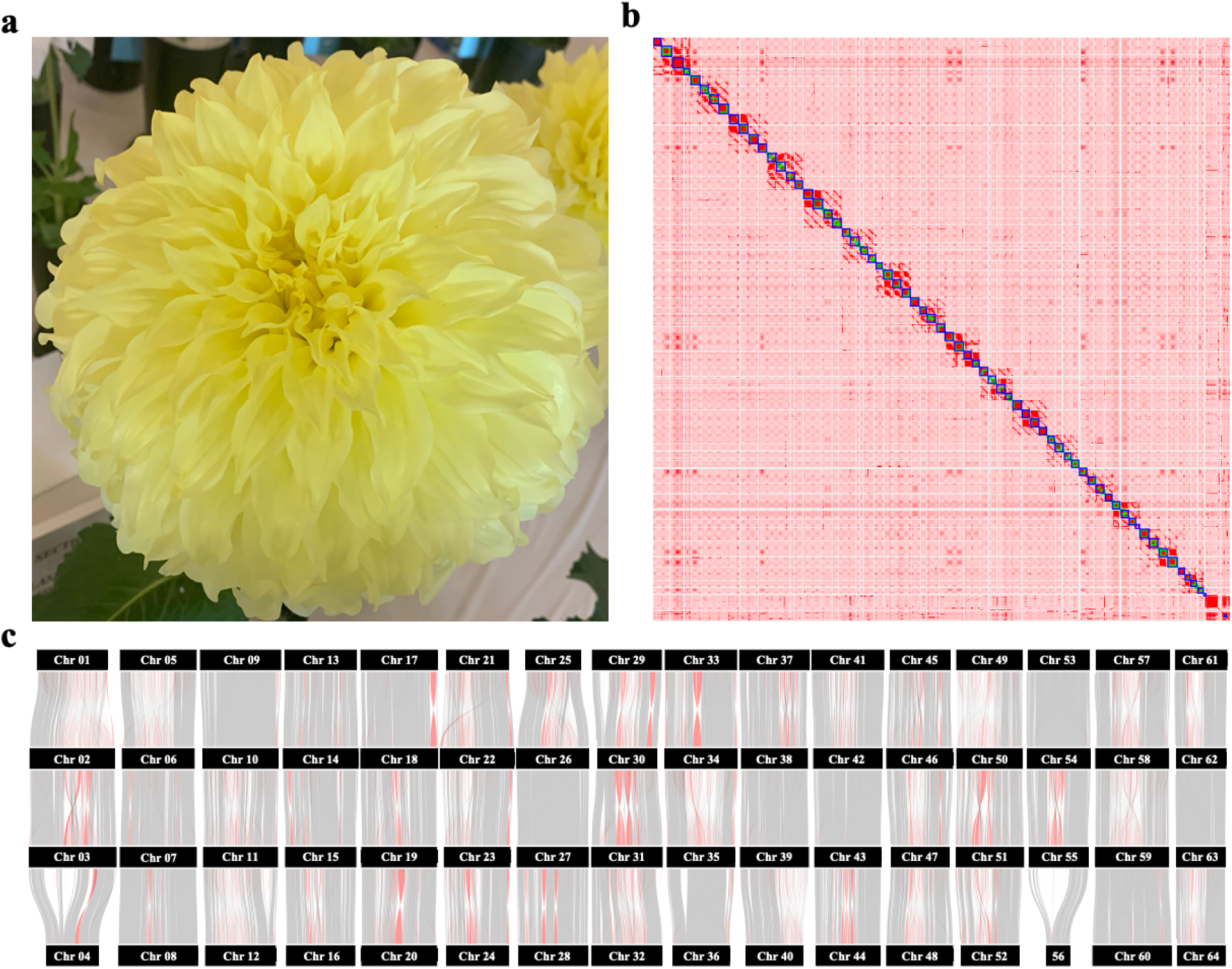
Hi-C maps and within-genome synteny comparison. A) Bloom of *Dahlia variabilis* ‘Edna C’, NEED A SCALE BAR ON THE PHOTO B) Contact map of chromosomal contacts of *Dahlia variabilis* ‘Edna C’. C) Nucleotide synteny of *Dahlia variabilis* within homologous chromosome groups.

## Methods & Materials

### Sample Collection, Extraction, and Sequencing

Young leaves of *Dahlia variabilis* cultivar Edna C were collected and flash-frozen in liquid nitrogen for high molecular weight (HMW) DNA extractions and Dovetail Omni-C library preparation. For HMW DNA isolation, three grams of young leaf tissue were ground in liquid nitrogen using a mortar and pestle and used as input for a Takara NucleoBond HMW extraction, following the manufacturer’s standard protocol (Takara Bio, Kusatsu, Japan). PacBio sequencing libraries were produced using the SMRTbell Express Template Prep Kit 2.0 chemistry and sequenced on 8 flow cells using a PacBio Sequel-II instrument in CCS/HiFi mode (Pacific Biosciences, Menlo Park, CA, USA). One gram of frozen young leaf tissue was used as input for a Dovetail Omni-C library preparation following the manufacturer’s protocol (Dovetail Genomics, Scotts Valley, CA, USA) and sequenced on an Illumina NovaSeq 6000 with PE150 reads (Illumina, San Diego, CA, USA). For RNA isolations, 0.02-0.15 grams of tissue were collected from 10 distinct developmental stages (roots, stem, young leaf, shoot apical meristem, 3 stages of developing inflorescences, disc florets, ray florets, and eyes), used as input for Trizol RNA extraction, and cleaned with a Zymo RNA Clean and Concentrator-25 before being sent to Novogene (Novogene, Beijing, China) for directional mRNA-seq library preparation and sequencing on a NovaSeq X Plus with PE150 reads.

### Genome Assembly

Genome size and polyploidy estimations were conducted with the PacBio HiFi CCS reads using *k*-mer spectra generated by KMC v 3.2.1 (Kokot 2017) with default parameters, GenomeScope V2.0 and Smudgeplot v0.2.5 (Ranallo-Benavidez 2020). Raw PacBio HiFi CCS reads were assembled with hifiasm v19.9 (Cheng 2024) with Omni-C integration under default parameters. The pseudohaplotype assemblies were polished with the PacBio reads using racon v1.5.0 (Vaser 2017), and then contigs smaller than 50 kb were removed with BBtools reformat.sh (Bushnell 2015). The assemblies were screened for contaminants using FCS-GX v0.5.4-5-g5dfd516 (Astashyn 2024) before scaffolding. Omni-C reads were mapped to the polished assemblies with bwa mem v0.7.17 (Heng 2009) with flags “-5SP -t 40” and samtools v1.10 (Heng 2009) with the flags “view -S -h -b -F 2316”. Sorted .bam files were used as input with Pairtools v1.0.3 (Open2C 2024) under default parameters to filter for duplicate, unmapped, and uniquely mapping reads to each assembly. These uniquely-mapped reads were used as input for YaHS v1.1 (Zhou 2023) with default parameters to scaffold the assemblies. Chromosomal contact maps were generated with juicer v1.1, juicer_tools v1.9.9, and Juicebox v2.17.00 using default parameters (Durand 2016). Manual edits of the contact map were made in Juicebox before using juicer_tools to create the curated assemblies. After manual curation, an exploratory gene annotation was generated only to orient and identify homoeologous chromosome groups using Liftoff v1.6.3 (Shumate 2021) and AGAT v1.4.0 (Daint 2022) using agat_sp_extract_seqeuences.pl with flags “-p” from *D. pinnata* ‘Kelvin Floodlight’ (Wang 2024). The annotation files were visualized using GENESPACE v1.3.0 (Lovell 2022), ‘Edna C’ chromosomes were moved and oriented based on synteny to ‘Kelvin Floodlight’. Pseudohaplotype assemblies were combined and re-scaffolded with YaHS v1.1 (Zhou 2023) before being manually curated with Juicebox v2.17.00 (Durand 2016) to create a single assembly containing all 64 chromosomes. Telomere locations were checked using plot_contigs from GENESPACE v1.3.0 (Lovell 2022). The assembly was checked for collapsed regions using bwa mem v0.7.17 (Li 2009) with flags ”-t 80 -k 21 -T 21 -a -c 10”, samtools v1.10 (Li 2009) with the flags “view -S -h -b -F 2316” to create the HiFi reads .bam file. Samtools sort (Li 2009) was used to sort the HiFi bam file before being used in bedtools v2.28.0 (Quinlan 2010) makewindows with flags “-g, -w 100000, -s 10000” and bedtools v2.28.0 (Quinlan 2010) coverage with flags “-sorted, -b -mean” to create a read depth file with 100 kb sliding windows with a 10 kb step. The completeness of this assembly was checked with compleasm v0.2.7 (Huang 2023) with the eudicots_odb10 gene set, and merqury v1.3 (Rhie 2020).

### Chloroplast Assemblies and Phylogeny

A chloroplast genome for *Dahlia variabilis* ‘Edna C’ was assembled with OatK (Zhou 2024) using the PacBio CCS HiFi long reads. Chloroplasts were also assembled for *Bidens alba* and *Dahlia pinnata* ‘Kelvin Floodlight’ (Wang 2024) using the same methods. Additional chloroplasts were downloaded from NCBI for *Dahlia imperialis* (Duan 2023), *Dahlia pinnata* ’Chocolate’ (Duan 2023), *Bidens cosmoides* (Knope 2020), *Bidens alba* var. Radiata (Knope 2020), and *Helianthus annuus* ‘HA 383’ (Timme 2006; Supplemental Table 7). These chloroplasts were annotated and visualized using GeSeq and Chlorobox under default parameters (Tillich 2017). Chloroplasts were oriented to the ‘Edna C’ chloroplast assembly, then were rotated to start at *psbA*. Chloroplasts were then aligned using MAFFT v7.520 with flags “--auto” (Katoh 2013) and a species tree was constructed with IQTree2 v2.2.2.7 with flags “-m MFP -B 1000 -nt AUTO” (Minh 2020) using the best-fit model “TVM+F+I” assigned by IQTree2, and then visualized in R v4.4.1 (R Core Team 2024).

### Repeat and Gene Annotation

Repeats were annotated using RepeatModeler v2.0.6 (Flynn 2020) with flag “-nLTRStruct” across all 64 chromosomes individually. Transposable Element (TE) libraries for each chromosome were combined and redundant TEs were removed using CD-hit-est V4.8.1(Fu 2012) with flags “-c 0.8 -n 8 -T 40” and reclassified with RepeatClassifier v 2.0.6 (Flynn 2020) to create a consensus TE library for the entire genome. DeepTE (Yan 2020) was used to identify the remaining unknown repeats. This curated TE library was used to softmask the ‘Edna C’ genome with RepeatMasker v4.1.7 (Smit 2015), with the flags “-s, -a, -gff, -xsmall, and -lib”. The .out file from RepeatMasker was used as input along with CDS sequence for EDTA v2.2.2 (Ou 2019) with flags “--genome, --rmout, --cds, --anno 1, --threads 21, --force 1” to add structural evidence for repeat families. A repeat insertion landscape was created using the .align file and calcDivergenceFromAlign.pl. The locations of putative centromeres were identified using RepeatObserver V1 (Elphinstone 2025) and a self-identity heat map was produced using ModDotPlot v0.9.4 (Sweeten 2024) with default parameters. Two tandem repeat annotators were used to identify large tandem repeats throughout the genome: TRASH v2 (Wlodzimierz 2023) with default parameters and Tandem Repeat Finder v4.09.1 (TRF; Benson 1999) with flags “2 7 7 80 10 50 500 -f -d -m -h”. Large tandem monomers from TRASH were filtered for an average identify high score > 30 and appeared consecutive more than 1000 times in a row capturing highly repetitive monomers. Similarly, tandem repeats from TRF were filtered for more than 1000 consecutive copies with a percent identity of 95% or greater. The softmasked genome was annotated with BRAKER3 v3.0.8 with additional flags “--gff3--threads 40” (Gabriel 2024) with mRNA-seq derived from the 10 diverse tissue types described above, along with protein evidence from closely related *D. pinnata, Cosmos bipinnata, Bidens alba,* and the Viridiplantae proteins. Genes were then filtered for the longest isoform using AGAT v1.4.0 (Daint 2022) with agat_sp_keep_longest_isoform.pl, then a peptide file was created with agat_sp_extract_sequence.pl, all using default parameters. The gene and repeat landscape was visualized using a custom python script available at (https://github.com/ZAMeharg/PolyploidHistoryofCultivatedDahlia).

### Novel Centromere Identification

Repeat families in the putative centromeric regions of *Dahlia variabilis* ‘Edna C’ were extracted and used to generate a centromeric repeat fasta file. This fasta was used as input for blastn v2.16.0+ with the flags “-outfmt 6 qseqid sseqid pident length qlen slen qstart qend sstart send evalue bitscore" -evalue 1e-10-max_target_seqs 10 -num_threads 80” (Altschul 1990) to several other Asteraceae: *Dahlia pinnata* ‘Kelvin Floodlight’, *Bidens alba* (Wang 2024), and *Helianthus annuus* 494 r1.0 (Badouin 2017). BLAST results were filtered for regions with a percent identity of 100% and a query length greater than 10,000 bp to reduce the noise of simple repeats blasting in highly-repetitive genomes. The putative centromeres of the additional Asteraceae assemblies were identified with TRF (Benson 1999) using the same method used to identify putative centromeres in ‘Edna C’. The location of the blasted repeats and TRF-identified centromeres were plotted using karyoploteR v1.30.0 (Gel & Serra 2017). A complementary analysis was also performed by extracting the TRF identified putative centromeric regions from *B. alba* and *D. pinnata* ‘Kelvin Floodlight’ using bedtools v2.28.0 getfasta (Quinlan 2010) and using them as input for blastn v2.16.0+ with flags “-outfmt 6 qseqid sseqid pident length qlen slen qstart qend sstart send evalue bitscore" -evalue 1e-10 -max_target_seqs 10 -num_threads 80” (Altschul 1990) to find the location of these repeats in *D. variabilis* ‘Edna C’ chromosomes. Seqkit v2.8.2 was used to identify the exact location of the centromeric monomer identified using TRASH V2 from ‘Edna C’ cross *D. pinnata* ‘Kelvin Floodlight’, *B. alba*, and *Helinathus annuus* 494 r1.0.

### Chromosome Partitioning

To partition the polyploid ‘Edna C’ assembly into putative subgenomes, Meryl v 1.4.1 with flags “count, k=25” (Rhie 2020) was used to *k*-merize all 64 chromosomes into *k*-mers of length 25. *K*-mers with an abundance of 10x per chromosome were identified with meryl “print greater-than 10” and combined to form a *k*-mer table that was imported to RStudio for filtering and normalization following a modified workflow (Session & Rokshar. 2023; https://github.com/amsession/Kmer-based-Subgenome-Mapping). *K*-mers were filtered and normalized in R v4.6.0 (R Core Team 2024) following a modified version from Sessions clustering script *k*-mer workshop for *k*-mers that are present at a 5 fold-change in one chromosome within a homoeologous group, and present on all chromosomes. This *k*-mer list was used to hierarchically cluster chromosomes and produce a heatmap of *k*-mer abundance. *K-*mers were also used to conduct a PCA using FactoMineR v2.11(Le 2008) to visualize chromosome clusters. Clusters were assigned to each chromosome based on the PCA results. Cluster-specific *k*-mers were identified and clustered into a PCA of *k*-mers and colored according to their respective cluster. The density of these *k*-mers were plotted onto each homoeologous chromosome group using blastn v2.16.0+ with flags “-evalue 10000 -word_size 25” to get exact matching of the cluster-specific *k*-mers and plotted with karyoplotter v1.30.0 kpPlotDensity with a 1,000,000 window size (Altschul 1990 et al; Gel & Serra 2017). Homoeologous chromosome relatedness was examined using single-copy gene trees from chromosomes and their orthologous chromosome in sunflower as an outgroup by Orthofinder v3.0.1b1 “-t 30, -a 30, -X” (Emms 2019) aligned with MAFFT v7.520 with flags “--auto” (Katoh 2013) and constructed with IQTree2 v2.2.2.7 with flags “-m MFP -B 1000 -nt AUTO --root-seq” (Minh 2020) where each homologous group was rooted to its syntenic chromosomes in sunflower, and combined into a coalescent tree using ASTRAL v5.7.8 (Zhang 2020) with default parameters, and visualized in R v4.4.1 (R Core Team 2024) using ggtree (Yu et al. 2017).

## Results and Discussion

### A tetraploid genome assembly for Dahlia variabilis

238 Gb of PacBio HiFi long-reads (∼32X tetraploid genome), and 102.4 Gb of Dovetail Omni-C short-reads (65,324,029 pairs or ∼13x) were generated to produce a *de novo* genome assembly for *D. variabilis* ‘Edna C’. *k*-mer based genome size estimations placed ‘Edna C’ at 1.3 Gbp per haplotype, with 6.3% heterozygosity (Supplemental Figure 1a). Polyploid estimation predicted ‘Edna C’ as a tetraploid with approximately 64% of *k*-mer pairs exhibiting an AAAB relationship pattern (Supplemental Figure 1b). Based on these estimations, we predicted the total genome size is roughly 5.2 Gb. Previous flow cytometry on other cultivated dahlias estimate the genome size between 8.0-9.6 Gb (Garnatje 2011, Lawrence 1929). This 2.8-4.4 Gb discrepancy between flow cytometry and *k*-mer based genome size estimates may be attributed to the polyploidy type. To account for these challenges, we explored several assembly routes. We assembled ‘Edna C’ into primary unitigs, primary contigs, and allowed hifiasm to partition the contigs into two pseudohaplotypes using the Omni-C reads. All of these assemblies were compared for base assembly statistics: total size, number of contigs, and contig N50 (Supplemental Table 1). While the primary unitigs assembly was within the estimated genome size range based on flow cytometry, the high number of unitigs (N=54,814) and the poor unitig N50 (666,483 bp) made downstream processing difficult. The contiguity issue of the primary unitig assembly was addressed with the primary contig assembly (N=3,606), but the genome size was roughly 2 Gb smaller (5,930,251,647 bp) than the primary unitig assembly. However, the contiguity and estimated genome size were in better concordance with previous reports when combining pseudohaplotype 1 (N=3,678 contigs, 5,891,493,068 bp) and pseudohaplotype 2 (N=1,103 contigs, 1,912,966,064 bp, Supplemental Table 1, Supplemental Figure 2). Combining both pseudohaplotypes generated the final assembly of 64 chromosomes with 16 groups of four chromosomes, leading to a tetraploid genome size of 7.73 Gb (Figure 1b, Table 1).

**Table 1:** Assembly Statistics for *Dahlia variabilis* ‘Edna C’.

| Assembly Statistics | Dahlia_variabilis_Edna_C.V1.Chr.fa |  |
| --- | --- | --- |
| Total Assembly Size (bp) | 7,728,871,057 |  |
| # of Contigs | 3,960 |  |
| Contig N50 (bp) | 21,743,008 |  |
| Longest Contig (bp) | 109,102,820 |  |
| # of Scaffolds | 1,942 |  |
| Scaffold N50 (bp) | 119,685,865 |  |
| Longest Scaffold (bp) | 149,452,150 |  |
| K-mer Completeness (%) | 97.0% |  |
| QV score | 48.3495 |  |
| BUSCO completeness (%) | 2280 | 98.02% |
| Single | 53 | 2.28% |
| Duplicated | 2227 | 95.74% |
| Fragmented | 15 | 0.64% |
| Incomplete | 0 | 0.00% |
| Missing | 31 | 1.33% |

We identified assembly collapses throughout the genome using several approaches, including visual identification during manual curation with Juicebox Assembly Tools, depth of Hifi reads mapped back to the assembly, and nucleotide synteny within homoeologous groups (Supplemental Figure 3). Altogether, 34 chromosomes had some level of collapse, totaling 136 Mbp (Figure 2, Supplemental Figure 4, Supplemental Table 2). The assembly BUSCO completeness scored 98% and the 64 chromosomes had a contig N50 of 21.7 Mb (Table 1, Supplemental Figure 5).

**Figure 2:**
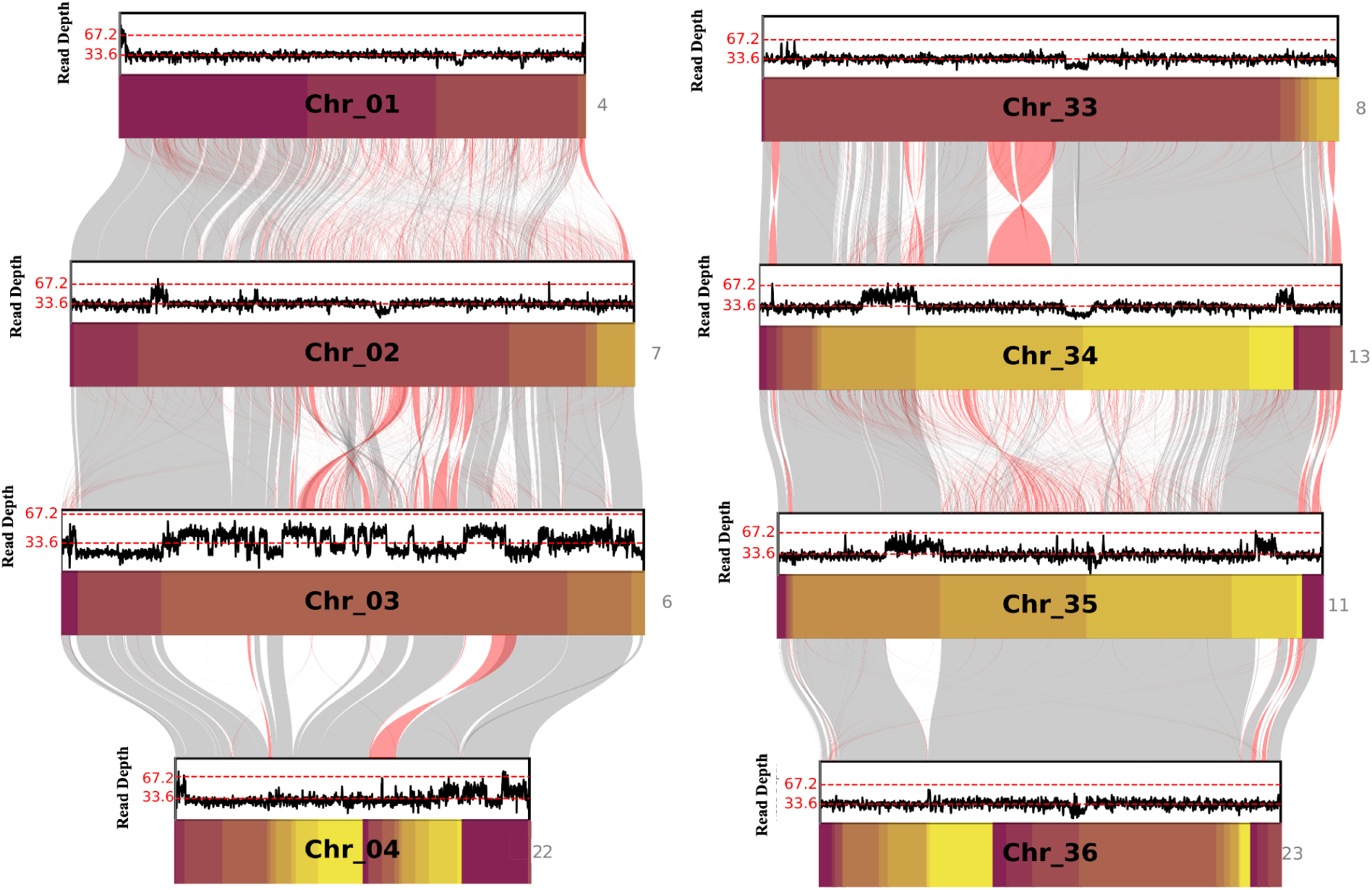
HiFi depth plots. A) A homologous group of ‘Edna C’ chromosomes where collapsed regions have been identified on Chromosome 3. B) A homologous group of ‘Edna C’ chromosomes where none of the chromosomes have indication of collapsed chromosomes.

*Dahlia variabilis* ‘Edna C’, like other Asteraceae genomes (Moore-Pollard 2025), has a high percentage of repetitive sequence at roughly 83.97% of the genome (Table 2). The repeat classifications indicated most were Long Terminal Repeats elements (LTR), 65.68%, 1.83% DNA transposons, 2.16% of simple repeats, and 12.96% Unknown. DeepTE further resolved the unknown repeats, adjusting the repeat percentages to 70% LTR, 9.67% DNA transposons, 2.16% simple repeats, and 1.10% unknown repeats (Supplemental Table 3). 141,996 protein-coding genes were annotated with BRAKER3, which is roughly 20,000 protein-coding genes fewer than found in *D. pinnata* ‘Kelvin Floodlight’, partially explained by the amount of collapse identified within the ‘Edna C’ assembly (Supplemental Figure 6, Table 2). Genes and repeats were layered onto synteny within homologous chromosome groups (Figure 3). Of the 34 chromosomes with predicted collapsed regions, nine of them contained collapses in the gene-dense arms of the chromosome, a feature which could explain the lower number of genes identified compared to ‘Kelvin Floodlight’. The remaining 25 chromosomes had various amounts of collapse in repeat-dense regions of their respective chromosomes.

**Figure 3:**
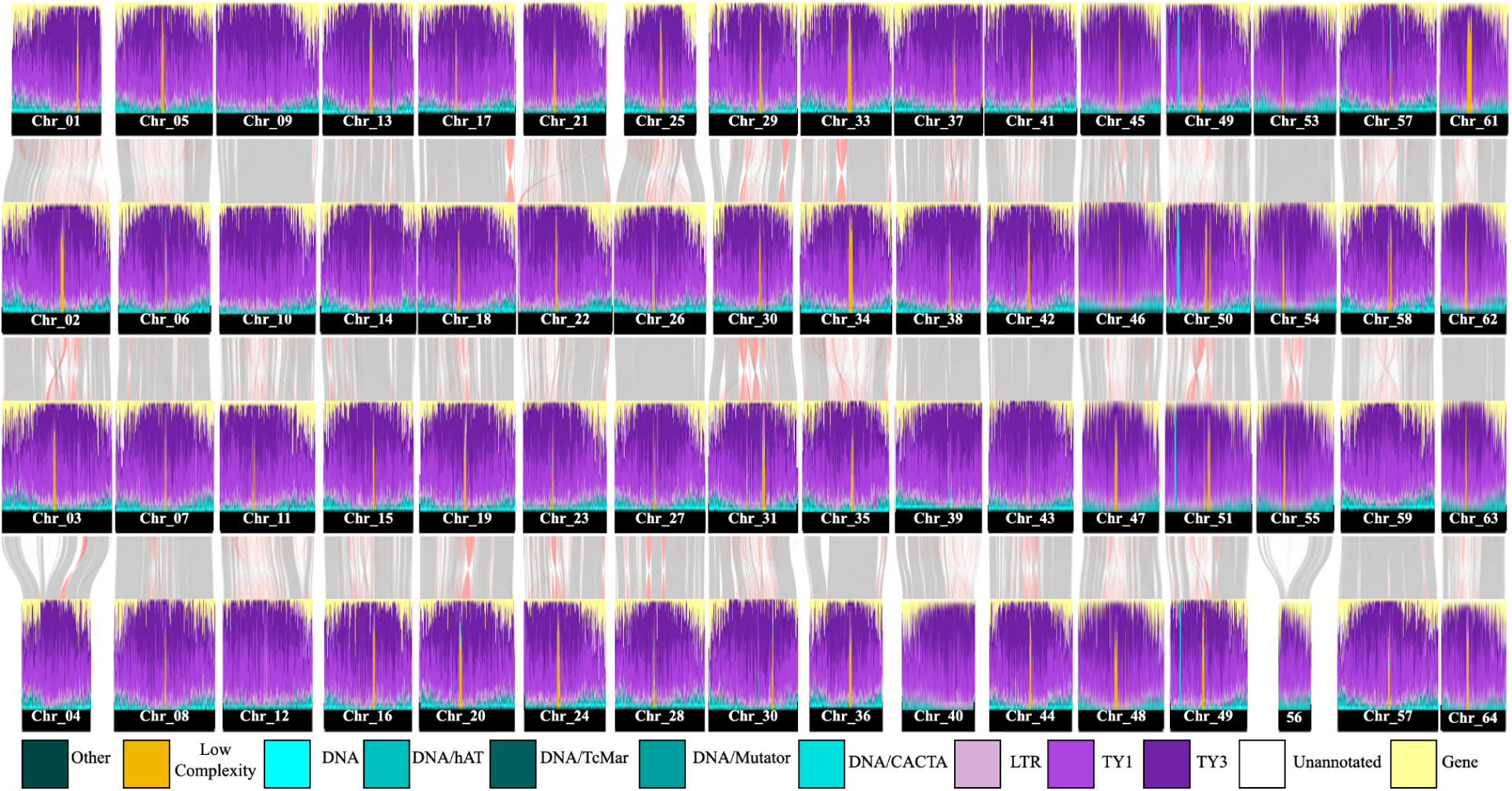
Gene and Repeat Landscape of Homoeologous Chromosome Groups of ‘Edna C’. A density plot of gene and repeat features across all 64 chromosomes. Genes are indicated in light yellow, retrotransposons are labeled in shade of purple, DNA transposons are labeled in cyan, and simple repeats are labeled as gold. Nucleotide synteny is also shown between chromosomes within homoeologous groups with grey indicating matching synteny and red indicating an inversion between chromosomes.

**Table 2:** Annotation Statistics.

| Annotation Statistics | Dahlia_variabilis_Edna_C.V1.Chr.fa |  |
| --- | --- | --- |
| Length of TE sequences (bp) | 6,480,829,186 (83.97%) |  |
| Number of protein-coding gene models |  | 141,996 |
| Number of protein-coding transcripts |  | 163,355 |
| BUSCO completeness of gene set (%) | 2,286 | 98.3% |
| Single | 16 | 0.7% |
| Duplicated | 2,270 | 97.6% |
| Fragmented | 7 | 0.3% |
| Incomplete | 0 | 0% |
| Missing | 33 | 1.4% |

### A highly repetitive genome reveals non-canonical predicted centromere motifs

We observed an abundance of low complexity repeats on 60 of the 64 chromosomes (Figure 3, Supplemental Figure 7). These regions were identified as putative centromeric regions through several complementary methods, including Shannon entropy, self-identification plots, repeat spectra heatmaps, and large tandem arrays (Supplemental Figure 8-9, Supplemental Table 3). Comparing the location of putative centromeric regions with the chromosome landscape, we identified a large number of simple or low-complexity repeats localized within the putative centromeric regions. In closely related *Helianthus*, centromere motifs have been found to be 187 bp (Quénet et al. 2012). In contrast to those, we detected a 20 bp motif, (ATGGCGTGGTATGGCGTGGT) present on 60 of the 64 chromosomes (Supplemental Table 4, Supplemental Table 5, Supplemental Figure 8). The four chromosomes missing this 20 bp motif are chromosomes 4, 9, 10 and 56. Chromosomes 4 and 56 have large amounts of collapse in the metacentric regions of the chromosomes when compared to their homologous chromosomes.

To test the uniqueness of this 20 bp centromeric motif found in ‘Edna C’, we searched for this motif in the genomes of related species *Bidens alba* (Wang 2024) and *Helianthus annuus* 383 (Badouin 2017), as well as the *Dahlia pinnata* ‘Kelvin Floodlight’ (Wang 2024; Figure 4). The phylogenetic position of these additional genomes and ‘Edna C’ were identified and compared to existing chloroplast assemblies from the additional genomes, *D. pinnata* ‘Chocolate’, *Dahlia imperialis*, and *Bidens alba* var *radiata* (Figure 4). The centromere-associated motifs for these three genomes were compared to ‘Edna C’ centromeric motif using two approaches: a reciprocal BLAST-based search where the respective motifs were filtered for a 90% identity to genome, and exact matching of the 20-bp motif. These two approaches demonstrated that when partial matches were allowed, segments of the motif were identified throughout most of the chromosome but not in that species’ respective centromere region. Using the exact matches approach, only putative centromeric regions in ‘Edna C’ and ‘Kelvin Floodlight’ had hits for the ‘Edna C’ motif and were absent in the other genome, indicating ‘Edna C’ and ‘Kelvin Floodlight’ shared a cultivated dahlia-specific centromeric motif (Figure 4). To validate this, we used the same methods used to identify the motif in ‘Edna C’ with ‘Kelvin Floodlight, indicating the same 20 bp motif on all 64 chromosomes. We hypothesize that cultivated *Dahlias* share this 20 bp centromeric motif, and that it arose sometime in cultivated *Dahlia*’s history after the divergence of *Dahlia* from the rest of Coreopsideae. Determining if this motif is shared with other species in the *Dahlia* genus will require additional sequencing of wild species.

**Figure 4:**
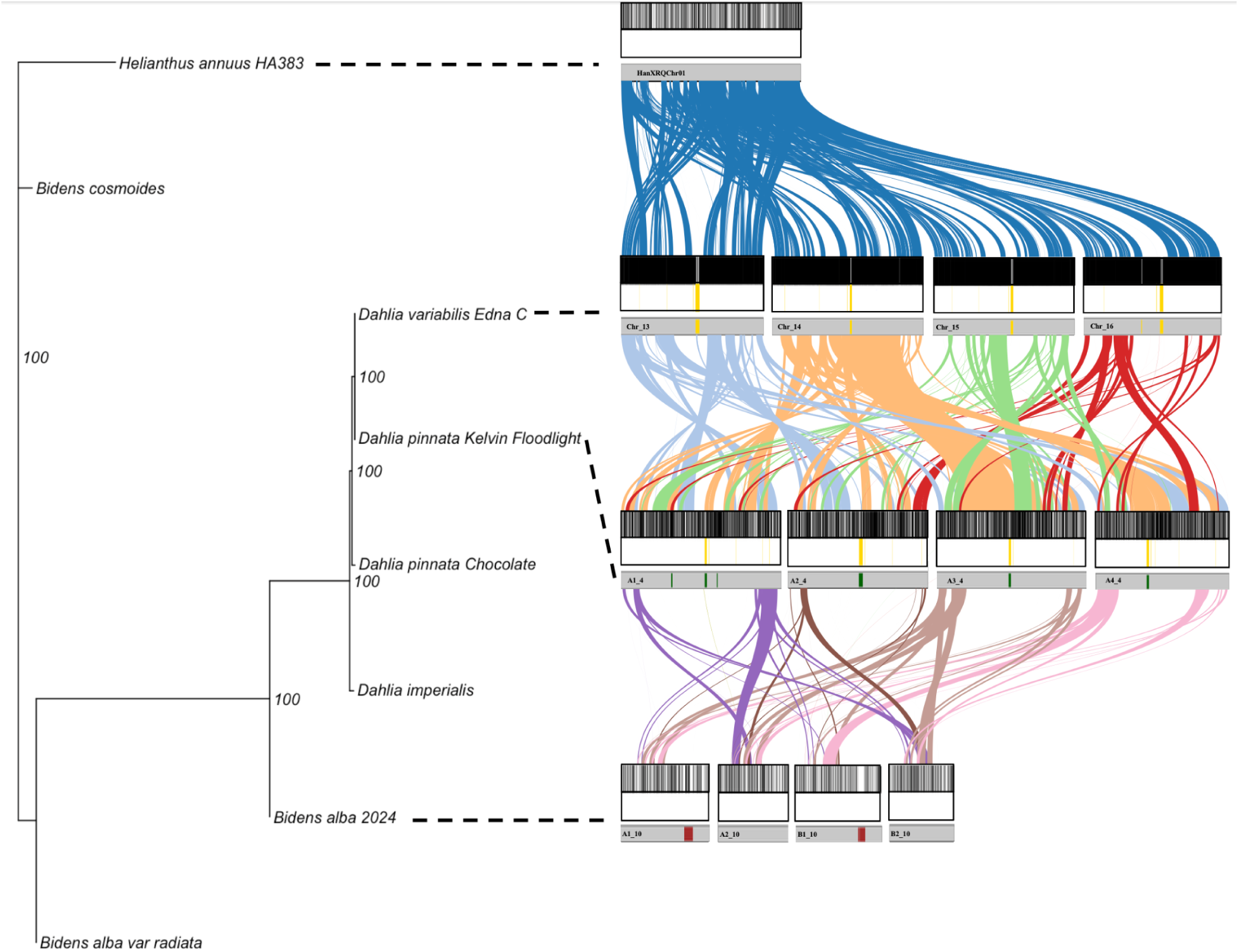
Novelty of ‘Edna C’ Centromere motif. A chloroplast species tree on the left to show the relatedness between *Helinathus, Bidens*, and *Dahlia*. On the right, a nucleotide synteny plot between orthologous chromosomes of *Helianthus, Dahlia variabilis ‘Edna C’, Dahlia pinnata ‘Kelvin Floodlight’,* and *Bidens alba*. The coordinates of the ‘Edna C’ centromeric motif are plotted above each chromosome, with those strictly matching directly above the chromosomes, and partially matching on top.

### K-mer based subgenome partitioning and gene tree analysis

Based on the tetraploid genome architecture of ‘Edna C’ and previous studies stating cultivated dahlias as a segmental allooctoploid (Schie 2014), we expected to find evidence of an allopolyploidy event within ‘Edna C’. Another study proposed that *Dahlia pinnata* was an autotetraploid because of the difficulties to distinguish subgenome specific *k*-mers (Wang et al. 2024). In order to compare the type of polyploidy in ‘Edna C’ to *D. pinnata* in the absence of known progenitor species, we used repetitive *k*-mer based clustering and gene tree analysis within homoeologous chromosome groups to characterize putative subgenome structure. We identified the subgenomes using transposable elements (TEs) that would be derived from the different progenitor species (Session et al. 2023). We excluded chromosomes that were less than 70% of the average chromosome size within their respective homologous group, which removed Chromosome 56 that was nearly 55% collapsed (Supplemental Table 8). With 63 of the 64 chromosomes, we extracted highly repetitive 25-mers found at least 10x per chromosome and normalized the count based on length of the chromosome. Because there is lineage-specific accumulation of TEs, the progenitor species likely accumulated these repetitive elements at different rates prior to the allopolyploidy event. We filtered the 25-mer list for 25-mers that exhibited a 5 fold change between any of the chromosomes within a homoeologous group. This identified 2,101 25-mers that were used to identify three distinct groupings of chromosomes, called Cluster-1, Cluster-2, and Cluster-3 with 12, 37, and 14 chromosomes respectively. These groupings show the same cluster patterns with a *k*-mer abundance heatmap (Figure 5a). Additionally, a PCA of the chromosomes shows the same pattern of the three clusters (Figure 5b).

**Figure 5:**
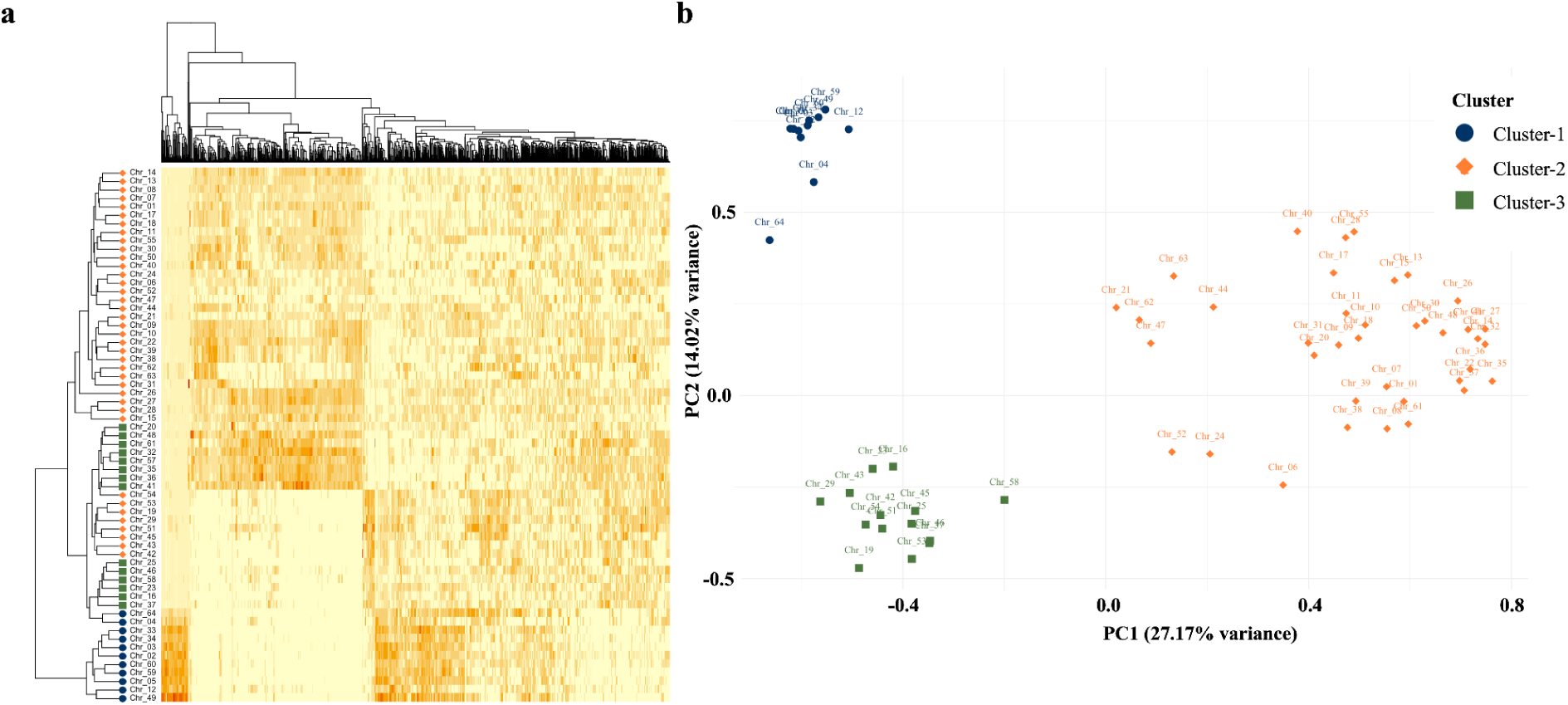
Chromosome Partitioning using Repetitive *k*-mers. a) A heatmap generated based on hierarchical clustering of highly repetitive *k*-mers showing the abundance of that *k*-mer on that particular chromosome. b) A principal component analysis of the repetitive *k*-mers on each chromosome to indicate how chromosomes cluster in relation to each other, the PCA shows three clusters (Cluster 1 as blue circles, Cluster 2 as orange diamonds, and Cluster 3 as green squares).

To determine the cluster-specific *k*-mer density on a given chromosome, we assigned each of the 2,102 *k*-mers to a cluster based on the highest average count across all the chromosomes in a given cluster. We identified 687 *k*-mers, 984 *k*-mers, and 431 *k*-mers for Clusters 1-3, respectively. The PCA of the *k*-mers showed the same cluster patterns found previously (Supplemental Figure 10). Chi-square goodness-of-fit tests were performed to determine whether the cluster-specific k-mers were distributed among chromosomes according to expectations based on chromosome length, which could suggest different species parentage. The null hypothesis of random chromosomal distribution was rejected for all three clusters (p < 2.2 × 10^-16^), indicating significant chromosomal enrichment of cluster-specific k-mers (Supplemental Table 9). To evaluate local variation in *k*-mer density, Kruskal-Wallis tests were conducted using counts within 1 Mb windows. Significant differences in *k*-mer density among chromosomes were detected for all three clusters (p < 2.2 × 10^-16^); box and whisker plots were generated to visualize the densities of cluster-specific *k*-mers across the whole genome, demonstrating that cluster-specific k-mers are not uniformly distributed across the genome (Supplemental Table 10; Supplemental Figures 11-13). The negative binomial model further demonstrated significant chromosome-specific differences in *k*-mer density, with several chromosomes exhibiting multiple-fold enrichment relative to the reference chromosome (Supplemental Tables 10-12). The location of these *k*-mers on the chromosomes were identified using the 1Mb windows previously generated (Figure 6a). Chromosomes had an abundance of *k*-mers with their assigned clusters, as well as portions of chromosomes that had greater abundance of other cluster *k*-mers usually in the arms of the chromosomes (Supplemental Figure 14). Chromosomes 12 and 64 are an example of this with both chromosomes clustering in Cluster-1 of the PCA (Figure 5a), but upon closer inspection of the *k*-mer density plots, Chr_12 has a greater density of Cluster-2 *k*-mers on the chromosome arms whereas Chr_64 has a greater density of Cluster-3 *k*-mers (Supplemental Figure 15).

**Figure 6:**
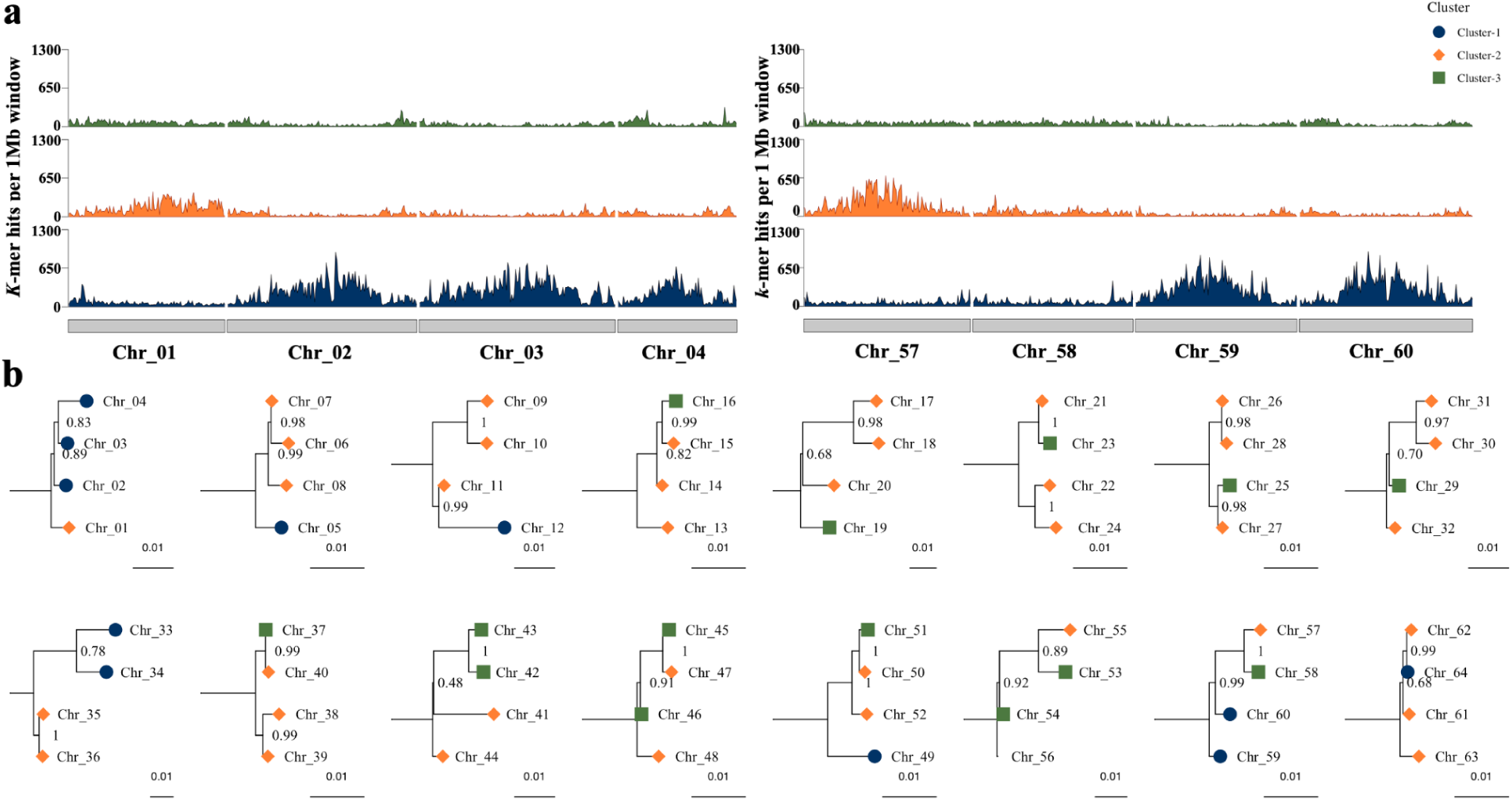
Cluster-specific *k*-mer Density and Cluster Relatedness. a) A *k*-mer density plot based on *k*-mers that passed through the enrichment pipeline and correspond to a specific cluster. The plot above each chromosome corresponds to the density of that cluster’s *k*-mers. Cluster-1 is on the bottom in blue, Cluster-2 is in the middle in orange, and Cluster-3 is on the top in green. b) Gene trees generated using single-copy genes from each homoeologous group and its orthologous chromosome in *H. annuus*. Clustering information is attached at the tip of each chromosome indicating the relatedness between homoeologous chromosomes.

After identification of three clusters of chromosomes, we hypothesize that *Dahlia variabilis* ‘Edna C’ arose from a combination of an auto- and allo-polyploidy events given the uneven distribution of chromosomes within clusters, unlike the previous assembly of *Dahlia pinnata* ‘Kelvin Floodlight’ that was identified as a strict autotetraploid. To address this, we investigated if the clustering in ‘Edna C’ is isolated to just ‘Edna C’ or if it is shared between both assembled cultivated dahlia genomes. Using the same *k*-mer clustering approach on *D. pinnata* ‘Kelvin Floodlight’ we identified two clusters based on a the *k*-mer abundance heatmap and clustering within the PCA of 1037 *k*-mers, with 15 and 49 chromosomes in Cluster-1 and Cluster-2 respectively (Supplemental Figure 16). Cluster-specific *k*-mer density supported this clustering, and followed a similar pattern as with ‘Edna C’. Chromosomes with a high density of Cluster-1 *k*-mers had a low density of Cluster-2 *k*-mers and vice versa (Supplemental Figure 17). To see if the identified clusters were derived from the same progenitor species in both ‘Edna C’ and ‘Kelvin Floodlight’ we plotted the density for each cultivar’s cluster-specific *k*-mers onto both cultivars indicating that ‘Edna C’ Cluster-1 and ‘Kelvin Floodlight’ Cluster-1 derived from the same progenitor species. Cluster-2 from ‘Edna C’ shared a high Cluster-2 density with ‘Kelvin Floodlight’ indicating these two clusters likely arose from the same progenitor species (Supplemental Figure 18). Cluster-3 was unique to ‘Edna C’ but both the *k*-mer density and abundance heatmap showed that Cluster-3 chromosomes still contained k-mers from the other two clusters at a lower abundance. This leads to the hypothesis that Cluster-3 may be derived from homoeologous exchange between Cluster-1 and Cluster-2, or is derived from a hybrid of the Cluster-1 progenitor and another progenitor not found in the ancestry of ‘Kelvin Floodlight’.

To further understand putative subgenome structure of cultivated *Dahlia variabilis* ‘Edna C’, we conducted a gene tree analysis by extracting each chromosome, generating pseudogenomes, and rooting with the syntenic chromosome of *H. annuus*, allowing us to investigate “single-copy” genes in a polyploid organism. We generated 16 trees (one for each homoeologous group) to examine the relationships between chromosomes (Figure 6b). These trees showed that Cluster-1 chromosomes were typically placed as the outgroup to the rest of the chromosomes within the group, with the exception of groups 3, and 16. Groups 3 and 16 place a Cluster-1 chromosome within a bifurcating branch or deep within the tree. Additionally, *k*-mer density plots identified several chromosomes that switch between clusters (Supplemental Figure 16). The change in the Cluster identity on the gene dense regions of the chromosomes explains the difference in tree topology and repetitive *k*-mer clustering. Chromosomes 12 and 64 cluster within Cluster-1 based on the *k*-mers, but the tree topology nest them with chromosomes from different clusters reflecting potential homoeologous exchange between the different clusters in homoeologous groups.

### Placing WGD in cultivated Dahlia

To investigate the placement of the polyploidy events in *D. variabilis* ‘Edna C’s evolutionary history, we calculated Ka and Ks across the entire genome to create a Ks plot (Figure 7a). Cultivated dahlia underwent two polyploid events, one more recent at a Ks value of 0.018, and another more distant at approximately 0.39. Between this Ks plot and previous literature, we hypothesize that the more distant whole genome duplication is likely an allopolyploidy event as a result of a hybridization between *D. coccinea* and *D. sorensenii* based on previous biochemical studies of the *Dahlia* (Ginannasi 1975). Further, the more recent genome duplication was caused by a hybridization event; we would expect to have observed a 2:2 relationship within all homoeologous groups. To test if this ancient allopolyploidy event occurred just within *Dahlia*, we used proteins from ‘Edna C’ and *Cosmos bipinnata* to generate an Edna C-*Cosmos* Ks plot to approximate the divergence between the two genera (Figure 7b). Based on the peak of that Ks curve (Ks ≈ 0.325) we were unable to determine if the whole genome duplication occurred before or after the divergence between *Dahlia* and *Cosmos*. Looking at the divergence of repeats throughout the genome, we hypothesize that divergence between the two genera occurred before the whole genome duplication, because we observed an increase in the divergence of the low-complexity centromeric motif unique to cultivated dahlia that started after the whole genome duplication event. If the divergence between *Dahlia* and *Cosmos* occurred during this proliferation of centromeric motifs, we would expect to find remnants of the low-complexity motif in the *Cosmos* genome yet we found no evidence of it in the genome. Additional whole genome assemblies will be necessary to fully untangle this complex evolutionary history.

**Figure 7:**
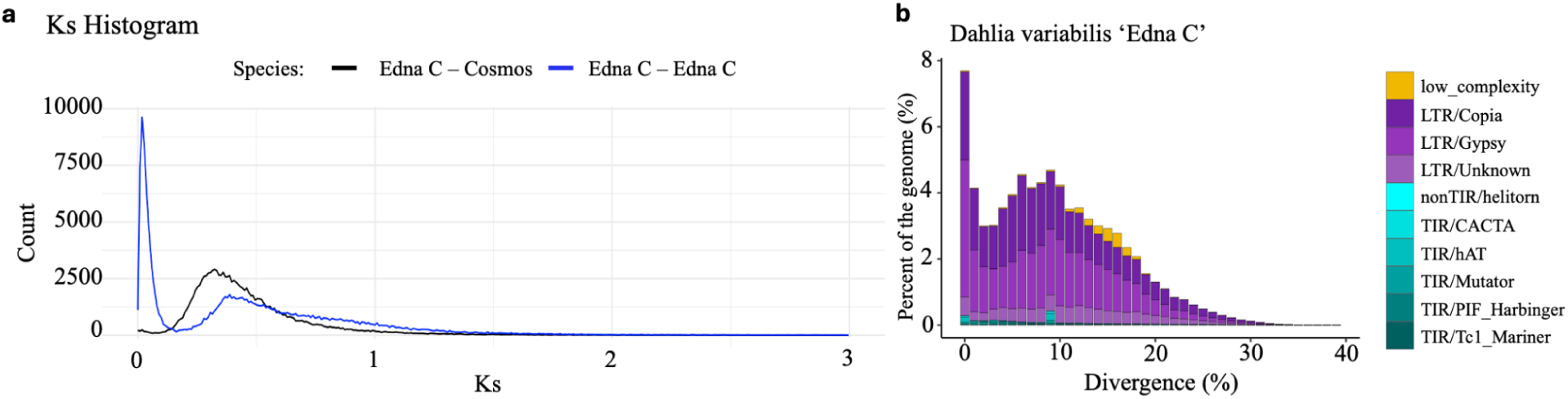
Genome duplication and repeat divergence in *Dahlia variabilis*. a) Ks histogram generated for ‘Edna C’, identified two peaks at roughly 0.03 and 0.39 in blue. A species divergence Ks distribution was generated to identify where *Dahlia* and *Cosmos* diverged from each other (at approximately 0.33). b) A repeat divergence plot showing percent divergence on the x-axis and percent of genome on the y-axis. Colors indicate the type of repeat family that diverged at that given percentage.

## Conclusion

This study presents the first chromosome-scale tetraploid genome assembly for *Dahlia variabilis* ‘Edna C’, providing an additional high-quality genome in the *Dahlia* genus. The ‘Edna C’ assembly is a repeat-rich genome that is predominantly LTR retrotransposons. Analysis of these repeat-rich regions identified a 20 bp tandem repeat that is strongly associated with the putative centromeric regions on nearly all chromosomes. Comparative analyses found this motif is shared between both our *D. variabilis* and *D. pinnata* but is absent from closely related genera – *H. annuus*, *C. bipinnata*, *B. alba*, suggesting this motif evolved after the divergence from the rest of the Coreopsideae tribe.

Investigation into the genome evolution provided evidence that cultivated dahlia does not fit a strict auto- or allotetraploid model. Repetitive *k*-mer clustering and single-copy gene tree analyses identified multiple clusters of chromosomes displaying potential signs of homoeologous exchange. These findings suggest that modern cultivars likely originated through a complex evolutionary history involving both allopolyploidy and autopolyploidy events, along with substantial genomic reshuffling through centuries of breeding and introgressions. The observation of a third chromosome cluster unique to ‘Edna C’ indicates that different cultivars may process distinct evolutionary histories contributing to the immense floral diversity within cultivated dahlia.

Two genome duplication events were identified, including an older duplication that was likely an ancestral allopolyploidization event near the divergence of *Dahlia* from the rest of the tribe. Second, there is a much more recent autopolyploid duplication event that likely contributed to the formation of modern cultivated dahlias. This analysis paired with the identification of a *Dahlia*-specific centromeric motif, provides a framework for the genome evolution and polyploid history of cultivated dahlia.

This genome assembly establishes an additional reference for Dahlia genomics and provides an insight into centromere evolution, polyploid genome evolution, and chromosome structure in a horticulturally-significant Asteraceae. Future sequencing of wild *Dahlia* species will be critical for identifying the progenitor lineages, testing the proposed evolutionary model, and facilitating molecular breeding and genetic improvement in cultivated dahlia.

## Acknowledgements

We thank the members of the American Dahlia Society for their support in funding data generation and plants sequenced for this project. Additionally we thank the members of the Kathy L. Chan Greenhouse staff at HudsonAlpha Institute for Biotechnology for their support in growing and maintaining the dahlia cultivars during the duration of this project.

## Data Availability

The raw data used to generate assemblies and annotations, as well as assemliblies are deposited under NCBI BioProject PRJNA1500019. Custom scripts for figures and analyses are housed at https://github.com/ZAMeharg/PolyploidHistoryofCultivatedDahlia.

## Funding

Z.M. and A.H. are funded through a generous donation from members of the American Dahlia Society. A.H. is funded by National Science Foundation IOS-PGRP CAREER Award #2239530.

## Conflicts of Interest

A.H. is a co-founder and board member of Veil Genomics.

